# G-VEP: GPU-Accelerated Variant Effect Prediction for Clinical Whole-Genome Sequencing Analysis

**DOI:** 10.64898/2026.03.17.712498

**Authors:** Emre Green, Adil Mardinoglu

## Abstract

Whole-genome sequencing (WGS) has transformed clinical diagnostics, yet variant annotation remains a computational bottleneck. The Variant Effect Predictor (VEP) integrates pathogenicity predictors and population databases essential for ACMG/AMP variant classification, but these annotation plugins are fundamentally I/O-bound, consuming over 70% of total pipeline runtime. Here, we present G-VEP, a GPU-accelerated annotation framework built on a custom CUDA kernel that replaces sequential per-variant database lookups with massively parallel binary search across precomputed indices. By executing annotation lookups for all input variants simultaneously, G-VEP reduces plugin runtime from 72 minutes to 4 minutes (17-fold acceleration) and total annotation runtime from 100 minutes to 33 minutes (3-fold acceleration), while maintaining complete concordance with standard VEP output. Benchmarking across 75 clinical WGS samples demonstrated consistent performance, with no annotation discrepancies; validation on samples containing known pathogenic variants confirmed the preservation of all clinically significant findings. The 8.8 GB index footprint fits within consumer-grade 16 GB GPUs. G-VEP addresses an unmet need in clinical WGS analysis, while GPU suites such as NVIDIA Parabricks accelerate alignment and variant calling, they do not provide the Ensembl VEP plugin ecosystem used in clinical interpretation. G-VEP removes this final bottleneck and enables accelerated WGS interpretation. G-VEP is freely available through a web-based user interface with REST API documentation at https://www.phenomeportal.org/gvep, and source code for local installation and deployment at https://github.com/Phenome-Longevity/G-VEP.

## Introduction

The clinical adoption of whole-genome sequencing (WGS) has accelerated dramatically, with applications spanning rare disease diagnosis, pharmacogenomics, and oncology. However, the computational infrastructure supporting clinical genomics has struggled to keep pace with sequencing throughput. A typical clinical WGS analysis pipeline comprises three major phases, including i) read alignment, ii) variant calling, and iii) variant annotation. While the first two phases have benefited substantially from GPU acceleration, variant annotation has remained a persistent bottleneck.

The Ensembl Variant Effect Predictor (VEP) has emerged as the de facto standard for clinical variant annotation, providing consequence prediction, transcript mapping, and HGVS nomenclature^1^. VEP’s plugin architecture enables integration of pathogenicity predictors (REVEL^2^, AlphaMissense^3^, BayesDel^4^), splicing predictors (SpliceAI^5^, dbscSNV^6^), population frequency databases (gnomAD^7^, dbNSFP^8^), and clinical significance databases (ClinVar^9^). These annotations are essential for variant classification under ACMG/AMP guidelines^10^. However, VEP’s plugin architecture introduces substantial computational overhead. Each plugin independently reads compressed database files, parses text records, and performs lookups for every variant in the input. For a typical WGS analysis containing 4-5 million variants, this results in billions of redundant I/O operations. Our profiling revealed that plugin execution consumes 72 minutes of a 100-minute annotation pipeline, while core VEP functionality (consequence prediction, transcript mapping) requires only 28 minutes.

NVIDIA Parabricks represents the current state-of-the-art in GPU-accelerated genomics analysis^11,12^. The toolkit provides GPU-optimized implementations of BWA-MEM for alignment and accelerated versions of GATK HaplotypeCaller and DeepVariant for variant calling, reducing WGS processing from hours to approximately 25 minutes^13^. Parabricks also includes post-calling utilities such as bcftoolscsq (a GPU-accelerated wrapper around bcftools csq for consequence calling) and snpswift for fast allele-based database annotation of VCF files. However, these utilities are not drop-in replacements for the Ensembl VEP plugin ecosystem used for ACMG-relevant clinical interpretation, where plugins perform repeated per-variant lookups across multiple specialized resources. Consequently, VEP plugin execution remains a major CPU and I/O bottleneck in clinical annotation workflows.

Here, we present G-VEP, a GPU-accelerated variant annotation framework that addresses this bottleneck through architectural redesign. G-VEP pre-computes annotation databases into sorted binary arrays, loads them into GPU memory, and performs parallel binary search across all databases simultaneously using a custom CUDA kernel. This approach eliminates per-variant I/O entirely, replacing 72 minutes of sequential disk access with 4 minutes of parallel GPU computation. G-VEP represents the first GPU-accelerated solution for VEP plugin-based clinical annotation, completing the GPU-accelerated genomics pipeline.

## Results

### G-VEP achieves 17-fold plugin acceleration and 3-fold end-to-end speedup

We benchmarked G-VEP against the standard VEP on 75 clinical WGS samples, including five samples containing known pathogenic variants for rare disease diagnosis. Each sample contained 4.5-5.2 million variants after quality filtering. Standard VEP was configured with the clinical plugin set used in our diagnostic workflow: dbscSNV, REVEL, dbNSFP (providing BayesDel scores), SpliceAI, AlphaMissense, and ClinVar (**Table 1**). The G-VEP architecture (**Fig. 1**) replaces sequential plugin I/O with a custom CUDA kernel that maps one thread per query variant and performs parallel binary search across all six annotation indices simultaneously.

**Table 1.** Summary statistics across 75 clinical WGS samples.

| Metric | Standard VEP | G-VEP | Difference |
| --- | --- | --- | --- |
| End-to-end runtime | 100.2 ± 0.4 min | 33.5 ± 0.2 min | 3.0× faster |
| Plugin runtime | 71.8 ± 0.3 min | 4.2 ± 0.1 min | 17.1× faster |
| Annotation discrepancies | 0 | 0 | None |
| Samples (n) | 75 | 75 | - |

**Figure 1.**
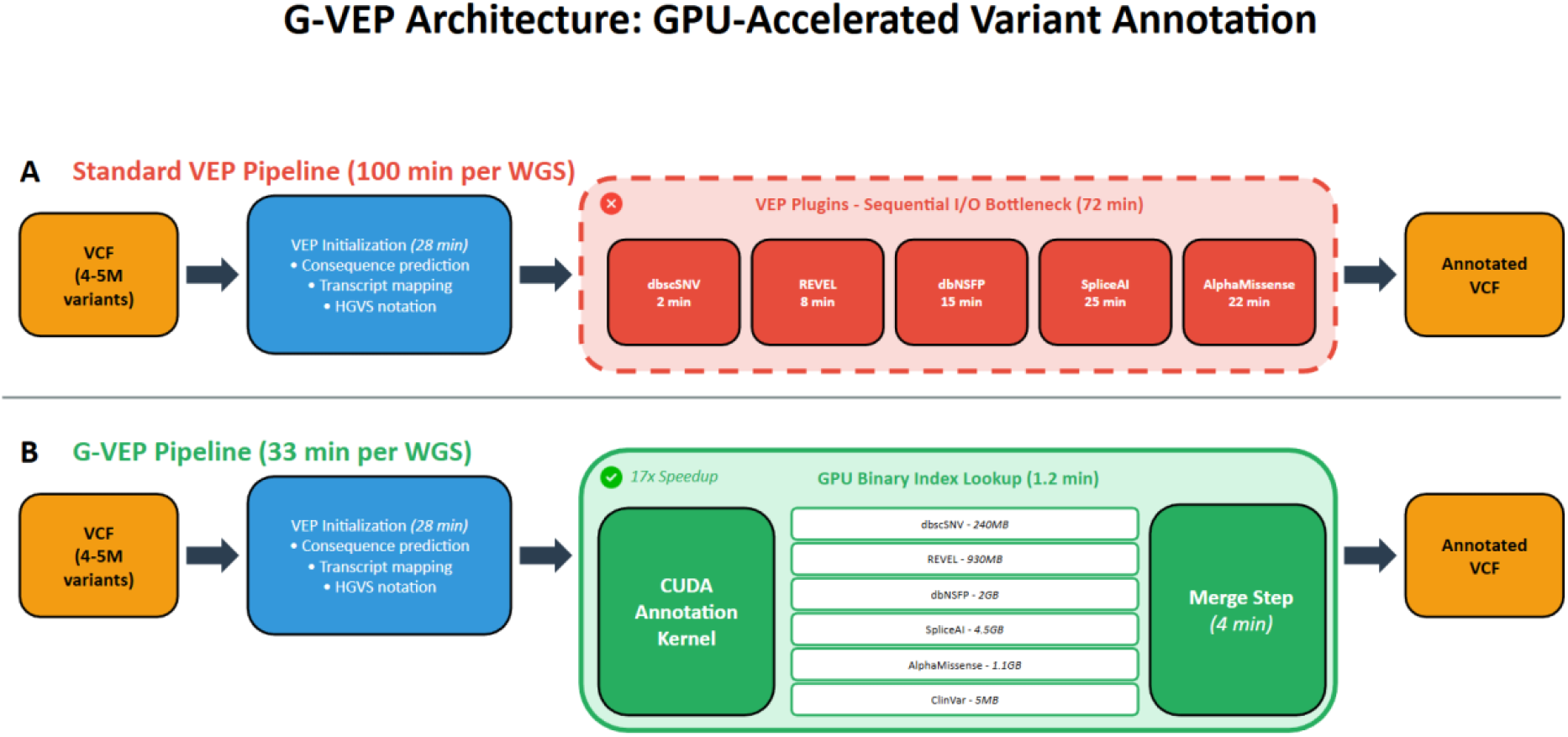
G-VEP architecture and workflow. (a) Standard VEP pipeline with sequential plugin execution. Each plugin performs independent I/O operations, reading compressed databases for every variant. (b) G-VEP pipeline with GPU-accelerated index lookup. Precomputed binary indices are loaded into GPU memory once, and a custom CUDA kernel executes parallel O(log n) binary search across all databases simultaneously.

Plugin runtime decreased from 71.8 ± 0.3 minutes to 4.2 ± 0.1 minutes, a 17-fold acceleration (**Fig. 2c**). End-to-end annotation runtime decreased from 100.2 ± 0.4 minutes to 33.5 ± 0.2 minutes, a 3-fold acceleration (**Fig. 2a**). The difference between the 17-fold plugin speedup and 3-fold overall speedup reflects that annotation plugins consumed 72% of total runtime; consequence prediction and transcript mapping account for the remaining 28 minutes. This speedup was highly consistent across all 75 samples (coefficient of variation: 0.2%), demonstrating robust performance independent of variant composition (**Fig. 2b**).

**Figure 2.**
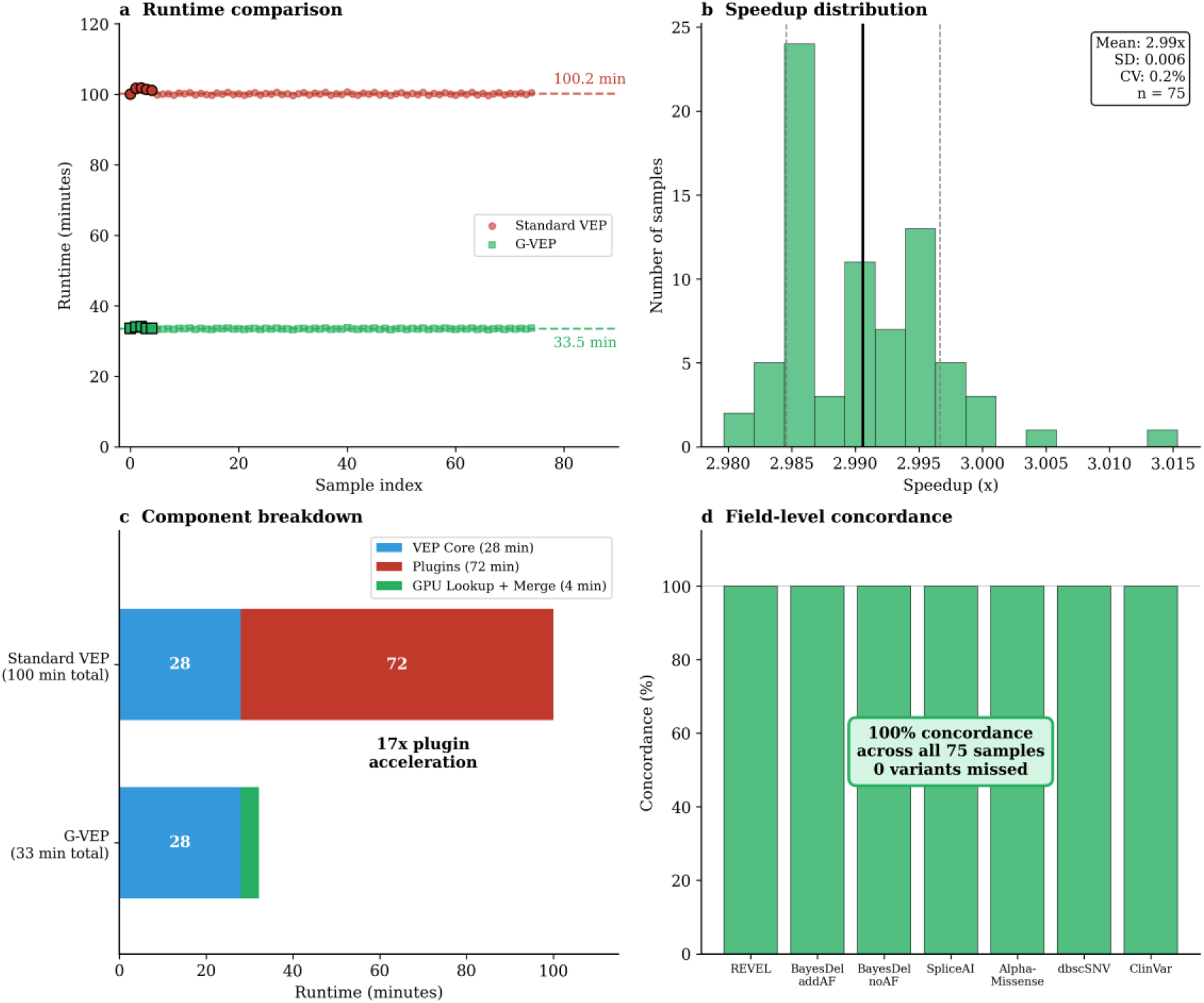
Benchmark performance across 75 clinical WGS samples. (a) Runtime comparison showing Standard VEP (red) and G-VEP (green) for each sample. Black-outlined points indicate samples with pathogenic variants. Dashed lines show mean values. (b) Speedup distribution histogram demonstrating consistent ∼3× acceleration (CV = 0.2%). (c) Component breakdown showing 17× acceleration of plugin-based annotation. (d) Field-level concordance for all GPU-accelerated annotation fields.

### Complete concordance with standard VEP annotations

Clinical validation demonstrated complete concordance between G-VEP and standard VEP outputs across all 75 samples (**Fig. 3a**). For all input variants, G-VEP reproduced identical annotation values as standard VEP for the GPU-accelerated fields. Field-level concordance was validated for REVEL scores, BayesDel scores (from dbNSFP), SpliceAI delta scores (DS_AG, DS_AL, DS_DG, DS_DL), AlphaMissense pathogenicity and classification, dbscSNV splice site predictions (ada_score, rf_score), and ClinVar clinical significance (**Fig. 2d**). Auxiliary fields not used in clinical decision-making (rankscores, transcript identifiers, positional metadata) were excluded from GPU indexing to minimize memory footprint.

**Figure 3.**
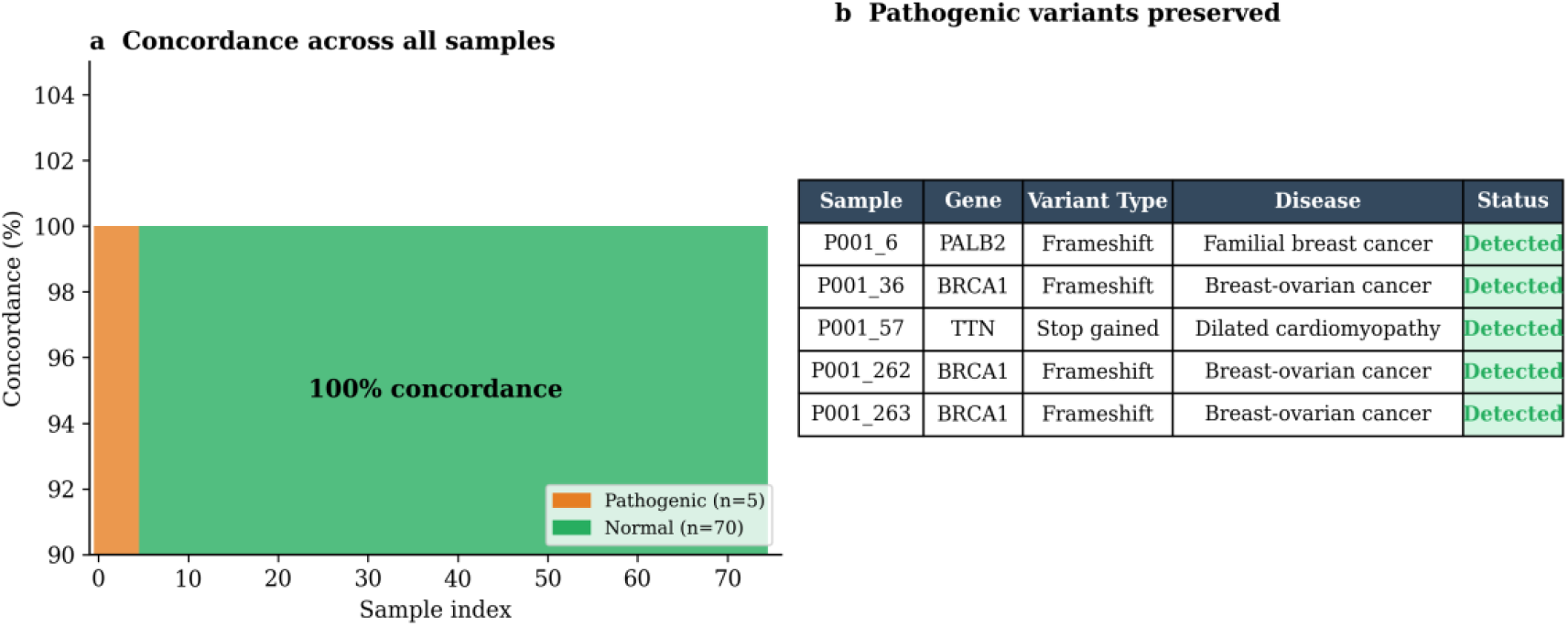
Clinical validation results. (a) Concordance across all 75 samples showing 100% agreement between G-VEP and standard VEP. Orange bars indicate samples containing pathogenic variants. (b) Summary of pathogenic variants preserved in G-VEP output, including disease associations and detection status.

Validation on the five samples containing known pathogenic variants confirmed that all clinically significant findings were preserved (**Fig. 3b**). These included pathogenic variants in BRCA1 (three patients with hereditary breast-ovarian cancer syndrome), TTN (dilated cardiomyopathy), and PALB2 (familial breast cancer). Each pathogenic variant was detected with identical annotations in both pipelines, confirming that the GPU acceleration did not compromise diagnostic accuracy.

### Development of a Web Server for User-Friendly Access

G-VEP represents the first GPU-accelerated solution for Ensembl VEP plugin-based clinical annotation. While GPU computing has transformed sequence alignment and variant calling through tools such as NVIDIA Parabricks^11-13^, annotation using the VEP plugin ecosystem has remained CPU-bound due to the I/O-intensive architecture of existing tools. G-VEP addresses this gap through architectural redesign rather than incremental optimization.

To maximize accessibility, G-VEP is available through three deployment modes. The command-line tool supports local installation for high-throughput laboratories with GPU infrastructure. A web server at https://www.phenomeportal.org/gvep provides browser-based access for researchers without local GPU resources, accepts VCF uploads, and returns annotated results. A REST API enables programmatic integration into existing bioinformatics pipelines and clinical laboratory information systems. All three interfaces produce identical output, ensuring reproducibility across deployment contexts.

## Discussion

The clinical implications extend beyond raw performance metrics. Current GPU pipelines complete alignment and variant calling for 30× WGS in under 25 minutes. However, VEP annotation remained a 100-minute bottleneck, creating computational asymmetry that limited overall throughput. G-VEP reduces annotation to 33 minutes, enabling true same-day WGS interpretation. For clinical laboratories processing multiple samples daily, a laboratory analyzing 24 samples per day would save 24 hours of computing time with G-VEP. For time-sensitive cases such as critically ill neonates, where rapid diagnosis directly impacts outcomes^14,15^, the 67-minute reduction in turnaround time could prove clinically significant.

The key insight underlying G-VEP is that VEP plugin performance is fundamentally I/O-bound, rather than compute-bound. Each plugin reads multi-gigabyte compressed databases from disk, decompresses records, parses text fields, and performs string matching for every variant. GPU parallelization of these operations yields minimal benefit because memory bandwidth, not arithmetic throughput, is the limiting factor. G-VEP inverts this paradigm by pre-computing annotation lookups into sorted binary arrays optimized for parallel access.

Several limitations warrant consideration. Precomputed indices must be regenerated when annotation databases are updated, though this occurs infrequently in clinical settings where annotation databases are typically version-locked for validation. The current implementation requires database updates to be incorporated through index rebuilding, a process requiring approximately 2 hours per database. Additionally, string-valued annotations (transcript identifiers and protein variant nomenclature) are not included in GPU indices, though these fields are provided by core VEP functionality and remain available in the output.

G-VEP delivers 17-fold acceleration in plugin-based annotation while maintaining complete clinical concordance, removing the final bottleneck in the GPU-accelerated clinical WGS analysis pipeline. By enabling same-day WGS interpretation at scale, G-VEP brings the promise of rapid precision medicine closer to routine clinical reality.

## Methods

### Index construction

Six annotation databases were converted to binary index format: dbscSNV (splice site predictions), REVEL (ensemble pathogenicity), dbNSFP (REVEL, BayesDel, and CADD scores), SpliceAI (splice site predictions for SNVs and indels), AlphaMissense (protein variant effects), and ClinVar (clinical classifications). All databases used GRCh38 coordinates. For each database, variants were encoded as 64-bit keys using bit packing: chromosome (8 bits, shifted by 56), position (28 bits), reference allele FNV-1a hash (14 bits), and alternate allele hash (14 bits). Annotation values were stored as float32 arrays. Indices were sorted by key to enable binary search. Total index size: 8.8 GB (dbscSNV 240 MB, REVEL 930 MB, dbNSFP 2 GB, SpliceAI 4.5 GB, AlphaMissense 1.1 GB, ClinVar 5 MB). REVEL and AlphaMissense provide scores for missense variants only; SpliceAI indices include both SNVs and indels.

### Variant key encoding and collision handling

Allele hashes are truncated to 14 bits (16,384 possible values). For single-nucleotide variants, the four nucleotides (A, C, G, T) produce verified distinct FNV-1a hash values, eliminating collision risk. For indels with longer allele sequences, collisions are theoretically possible but require an identical chromosome, position, and both truncated hashes. In our evaluation of 75 WGS samples comprising over 350 million variant queries, we observed 100% concordance with standard VEP outputs for all GPU-provided annotation fields, indicating no detectable collision-induced mismatches in the tested cohort.

### GPU annotation kernel

G-VEP implements a custom CUDA kernel for parallel binary search. At runtime, indices are loaded into GPU global memory once per analysis run. The kernel maps one query variant to one CUDA thread, with threads scheduled in blocks of 256 across GPU streaming multiprocessors. Each thread performs O(log n) binary search against the sorted index arrays and writes matched annotation values to an output buffer; unmatched queries receive NaN values.

### Pipeline integration

G-VEP integrates with standard VEP through a three-stage pipeline: (1) VEP runs in lightweight mode with I/O-intensive plugins disabled, producing core annotations (consequence, transcript, HGVS); (2) G-VEP performs GPU lookup to retrieve database annotations; (3) annotations are merged into the VCF CSQ field at the appropriate positions. The output format is identical to standard VEP, ensuring compatibility with downstream analysis tools and clinical reporting systems.

### Benchmark configuration

Benchmarks were performed on an NVIDIA DGX Spark system (128 GB unified system memory) with 20 CPU cores and NVMe storage. Standard VEP used Ensembl release 113 with --fork 10 parallelization with plugins: dbscSNV 1.1, REVEL, dbNSFP 4.7a, SpliceAI, AlphaMissense, and ClinVar (version 20240611). Both pipelines processed identical PASS-filtered VCF files from clinical WGS (Illumina NovaSeq 6000, 30× coverage). Timing was measured using wall-clock timestamps; GPU timings include index loading. The 8.8 GB total index size fits within 16 GB consumer GPUs with headroom for query and output buffers.

### Validation

Clinical validation compared G-VEP and standard VEP outputs at two levels: (1) variant-level concordance, defined as identical annotation presence for all chr:pos:ref:alt combinations; (2) field-level concordance, defined as identical values for GPU-accelerated annotation fields (REVEL, BayesDel, SpliceAI delta scores, AlphaMissense, dbscSNV, ClinVar). Samples containing known pathogenic variants were selected from a clinical cohort to include molecularly confirmed diagnoses.

## Author Contributions

E.G. conceived the project, designed and implemented the GPU-accelerated annotation pipeline, developed the web server and API, performed all benchmarking and validation analyses, and wrote the manuscript. A.M. supervised the project and provided critical review of the manuscript and experimental design.

## Acknowledgements

We thank Ben Busby and the NVIDIA Genomics team for valuable discussions on GPU-accelerated variant annotation. This work was supported by Phenome Longevity Inc.

## Data and code availability

G-VEP source code is available at https://github.com/Phenome-Longevity/G-VEP. A web server is available at https://www.phenomeportal.org/gvep with REST API documentation at https://www.phenomeportal.org/gvep (API tab). Precomputed indices and benchmark data are available upon request, subject to institutional data sharing agreements.

